# The record of Deinotheriidae from the Miocene of the Swiss Jura Mountains (Jura Canton, Switzerland)

**DOI:** 10.1101/2020.08.10.244061

**Authors:** Fanny Gagliardi, Olivier Maridet, Damien Becker

## Abstract

The Miocene sands of the Swiss Jura Mountains, long exploited in quarries for the construction industry, have yielded abundant fossil remains of large mammals. Among Deinotheriidae (Proboscidea), two species, *Prodeinotherium bavaricum* and *Deinotherium giganteum*, had previously been identified in the Delémont valley, but never described. A third species, *Deinotherium levius*, from the locality of Charmoille in Ajoie, is reported herein for the first time in Switzerland. These occurrences are dated from the middle to the late Miocene, correlating to the European Mammal biozones MN5 to MN9. The study is completed by a discussion on the palaeobiogeography of deinotheres at the European scale.

## Introduction

The order Proboscidea currently regroups large mammals whose common features include tusks and a long, muscular trunk. Within the superorder Afrotheria, its sister group is Sirenia (dugongs and manatees). Its extant representatives belong to the Elephantidae family with only three species of elephants living in Africa or Asia (*Loxodonta africana, Loxodonta cyclotis* and *Elephas maximus*). However, this order was much more diversified in the fossil record.

The proboscideans have an African origin hypothetically with the stem genus *Eritherium*, found in the early late Paleocene of Morocco (**Gheerbrant, 2009**), and indubitably with other primitive forms as the small-sized *Phosphatherium* and *Numidotherium* or the first large-sized proboscidean *Barytherium*. These primitive forms were only found in the late early Eocene, and the late Eocene and early Oligocene, respectively, of Africa (**Tassy, 1990**; **Sanders et al., 2010**). It should be noted that the relationship of *Eritherium* is unresolved. After **Gheerbrant et al. (2018)**, it is sister group to either both the Proboscidea and Sirenia or to all tethytherians. The gomphotheres (Gomphotheriidae), the mammutids (*Zygolophodon*) and the deinotheres (Deinotheriidae) are the earliest proboscideans found outside of Africa in the fossil record. Their occurrence in Europe is linked to the Proboscidean Datum Event (sensu **Tassy, 1990**) of the late early Miocene (ca. 19.5-17.5 Ma; **Göhlich, 1999**). This biogeographic event resulted from the counter clockwise rotation of Africa and Arabia plates leading to a collision with the Anatolian plate and the formation of a landbridge connecting Africa and Eurasia at the end of the early Miocene (**Rögl, 1999a,b**). This geographic change allowed remarkable terrestrial mammal exchanges including the gomphotheres and the deinotheres (e.g. **Göhlich, 1999**; **Sen, 2013**). Within the phylogeny of proboscideans (**Fig. 1**), deinotheres are included in a clade composed only of forms typically weighing more than 1000 kg (mega herbivores) together with Elephantiformes (*Phiomia, Mammut, Gomphotherium* and Elephantidae) of which they are the sister group (**Hutchinson et al., 2011**). The differentiation between deinotheres and Elephantiformes could have occurred as early as the end of the Eocene (e.g. **Delmer, 2009**). However, phylogenetic relationships within the Deinotheriidae family remain uncertain to this day.

**Figure 1.**
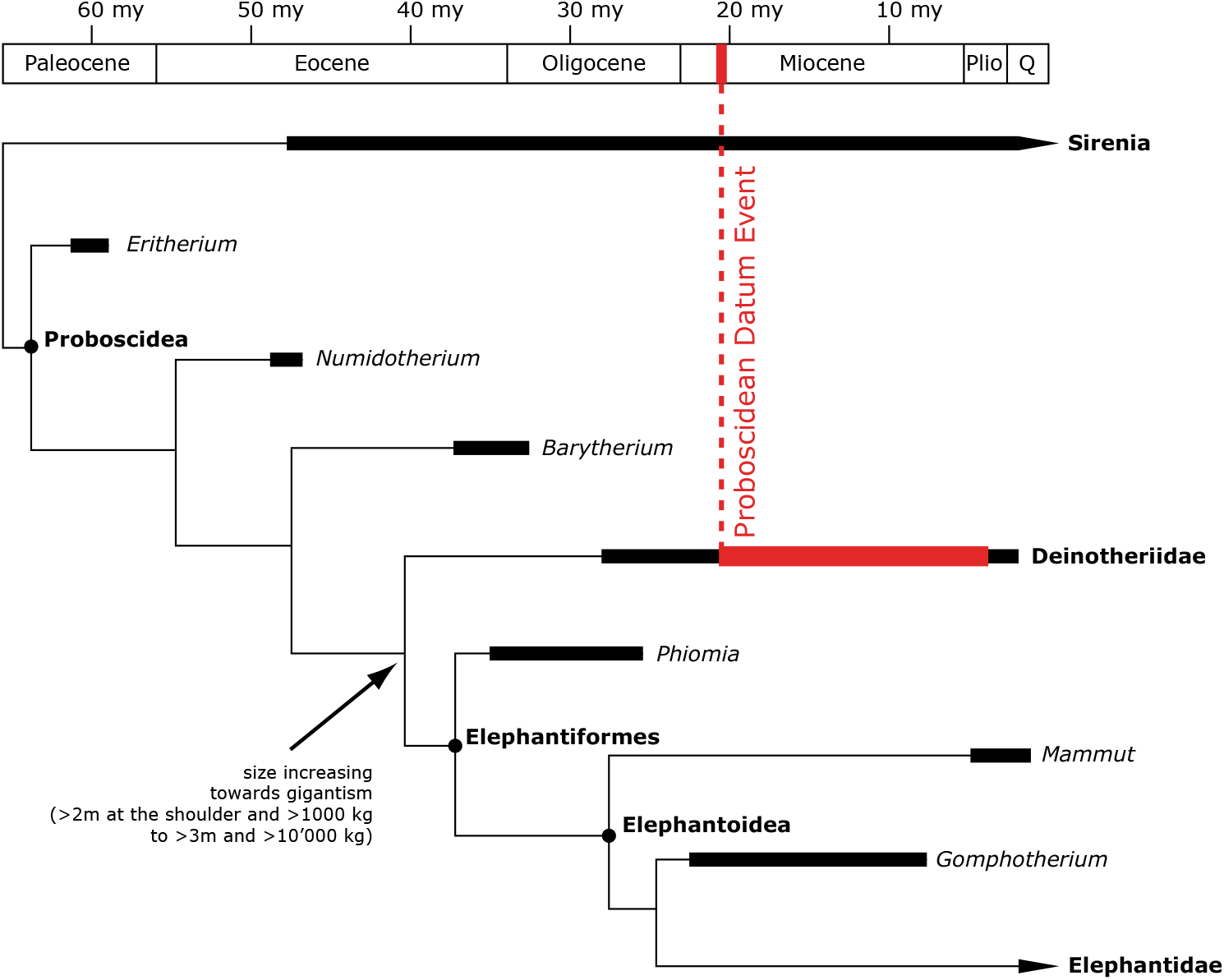
Simplified, stratigraphically calibrated, phylogeny of Proboscideans (modified from Hutchinson et al., 2011).

The oldest and most primitive deinothere, *Chilgatherium harrisi* **Sanders et al., 2004**, was discovered in Africa (Ethiopia) and is dated to the late Oligocene (**Sanders et al., 2004**). It disappeared slightly before the Miocene, probably replaced by *Prodeinotherium hobleyi* (**Andrews, 1911**) recorded in early Miocene of eastern Africa (**Harris, 1978**; **Pickford, 2003**; **Sanders et al., 2010**). After the Proboscidean Datum Event (ca. 19.5-17.5 Ma; late early Miocene), the distribution of the family extends to Asia with *Prodeinotherium pentapotamiae* (**Falconer, 1868**) discovered in Pakistan (**Welcomme et al., 1997**) and to Europe with *Prodeinotherium cuvieri* **Kaup, 1832** in Greece (MN3; specimens from Lesvos Island identified as *Prodeinotherium bavaricum* (**Meyer, 1831**) by **Koufos et al., 2003**, but corrected in *P. cuvieri* following criteria of **Ginsburg and Chevrier, 2001** and **Pickford and Pourabrishami, 2013**) as well as in France and Spain (MN4; **Azanza et al., 1993**; **Ginsburg and Chevrier, 2001**). *Prodeinotherium bavaricum* **Éhik, 1930** (= *P. hungaricum* after **Huttunen, 2002a**) is also recorded in the early Miocene in Hungary (**Éhik, 1930**; **Gasparik, 1993, 2001**). The last deinotheres are still present in Asia by the late Miocene with *Deinotherium giganteum* **Kaup, 1829**, *Deinotherium proavum* (**Eichwald, 1831**) (= *D. gigantissimum* after **Huttunen, 2002a**) and *Deinotherium indicum* **Falconer, 1845** (**Chaimanee et al., 2004**; **Rai, 2004**; **Singh et al., 2020**). In Africa, they persist with *Deinotherium bozasi* **Arambourg, 1934** until the early Pleistocene (**Harris, 1983**; **Harris et al., 1988**). In the fossil record of Europe, three species seemed to occur during the middle Miocene, although few evidences exist of an actual coexistence in fossil assemblages (e.g. **Duranthon et al., 2007**): *Prodeinotherium bavaricum, Deinotherium levius* **Jourdan, 1861** and *Deinotherium giganteum* (e.g. **Göhlich, 1999**; **Ginsburg and Chevrier, 2001**; **Pickford and Pourabrishami, 2013**). The latter survived until the end of the Vallesian, whereas during the Turolian *Deinotherium proavum* was the last representative of deinotheres in Europe (e.g. **Codrea et al., 2002**; **Kovachev and Nikolov, 2006**; **Boev and Spassov, 2009**; **Konidaris et al., 2017**).

From the Swiss Jura Mountains, **Bachmann (1875)** described a deinothere mandible in five fragments, discovered in the west of the Montchaibeux hill by Jean-Baptiste Greppin in 1869, which he referred to *Deinotherium bavaricum*. **Greppin (1867, 1870)** reported the presence of a lower molar of *Deinotherium giganteum*, discovered by the geologist and naturalist Peter Merian in 1858, in the forest of Bois de Raube of the Delémont valley. Deinotheres and gomphotheres were also found in Charmoille and successively reported by **Stehlin (1914), Schäfer (1961)** and **Kälin (1993)**. However, none of the deinothere remains from Charmoille have ever been described. Additionally, an isolated upper molar labelled *Deinotherium bavaricum* is housed in the Jurassica Museum collections. This specimen has never been reported before and its exact origin in the Delémont valley remains uncertain. This study focuses on the fossil remains of deinotheres discovered in the Swiss Jura Mountains in order to provide a complete description of the specimens and to update their identifications. A discussion on the distribution of deinotheres throughout the Miocene of Europe completes the article.

### Geographic, geologic and stratigraphic framework

The Jura Canton lies at the palaeogeographic junction between the Cenozoic tectonic and sedimentary provinces of the Upper Rhine Graben and the North Alpine Foreland Basin (**Sissingh, 2006**). The regional fluvio-lacustrine sediments of the Miocene Bois de Raube Formation (OSM; Obere Süsswassermolasse = Upper freshwater molasse), were deposited both in Delémont Basin (near Delémont) and in Ajoie area (near Porrentruy). According to **Kälin (1997)**, this formation is subdivided into three members differing by a markedly different heavy mineral spectrum and pebble content: a basal Montchaibeux Member (“RoteMergel und Dinotheriensande des Mont Chaibeux” of **Liniger, 1925**), a middle conglomeratic Bois de Raube Member (“Vogesenschotter des Bois de Raube” of **Liniger, 1925**) in Delémont Basin, and an upper Ajoie Member (“Hipparionsande von Charmoille” of **Liniger, 1925**). The formation covers the biochronological interval MN4 to MN9 (**Kälin, 1997**; **Choffat and Becker, 2017**; **Prieto et al., 2018**) and includes three historical localities yielding deinothere remains (**Greppin, 1867, 1870**; **Stehlin, 1914**; **Schäfer, 1961**; **Kälin, 1993**): Montchaibeux (MN5-6) in Rossemaison, Bois de Raube (MN7/8) in Develier, and Charmoille (MN9) in Ajoie (**Fig. 2**).

**Figure 2.**
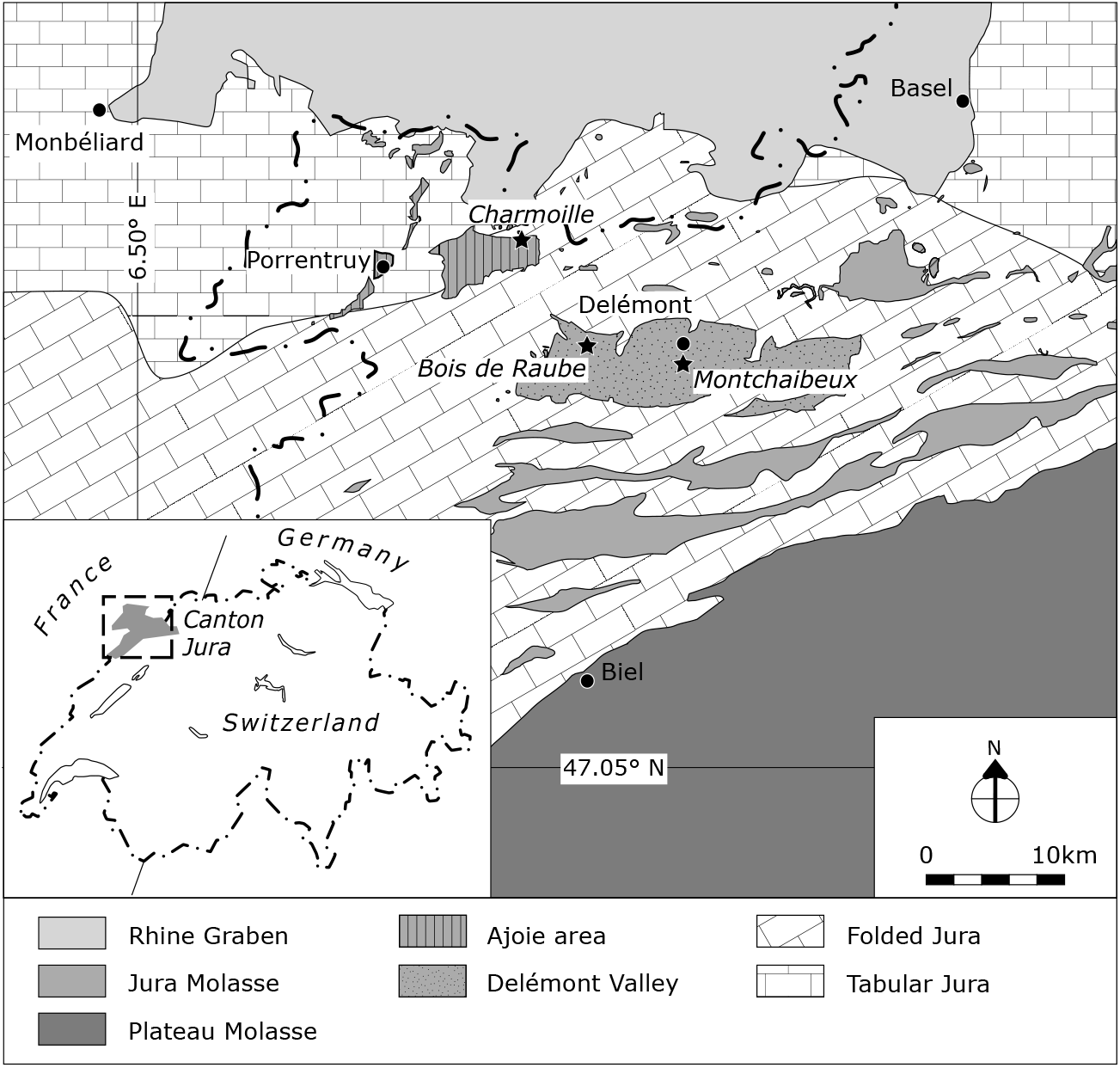
Geographic and geologic context of the Swiss Jura localities (Montchaibeux, Bois de Raube and Charmoille) with Deinotheriidae remains.

## Material and method

### Material

The studied material of Deinotheriidae, coming exclusively from the Swiss Jura Canton, includes:

1. The famous reconstituted mandible of *Prodeinotherium bavaricum* from the Montchaibeux locality (**Bachmann, 1875**). A copy of this mandible is housed in the collections of the Jurassica Museum whereas the original specimen is housed in the collections Natural History Museum of Bern;
2. A copy of the lower molar of *Deinotherium giganteum* from the Bois de Raube locality (**Greppin, 1867, 1870**), housed in the Jurassica Museum and whose original seems to be housed in the Jean-Baptiste Greppin collection of Strasbourg University;
3. The upper molar of the Jurassica Museum collection of *Prodeinotherium bavaricum* coming probably from the Delémont valley; and
4. The specimens of Deinotheriidae from Charmoille (**Stehlin, 1914**; **Schäfer, 1961**; **Kälin, 1993, 1997**; **Choffat and Becker, 2017**) which consist in some fragments of tusks from the Jurassica Museum collection and more complete dental specimens housed in the Museum of Natural History of Basel.

### Terminology and measurements

The dental terminology for Deinotheriidae mainly follows that of **Aiglstorfer et al. (2014)** and **Pickford and Pourabrishami (2013)**, and is illustrated in this paper (**Fig. 3**) for a better understanding of the character descriptions and discussions. The measurements written in the tables or in the text are given in millimetres (precision at 0.1 mm), those in brackets are estimated.

**Figure 3.**
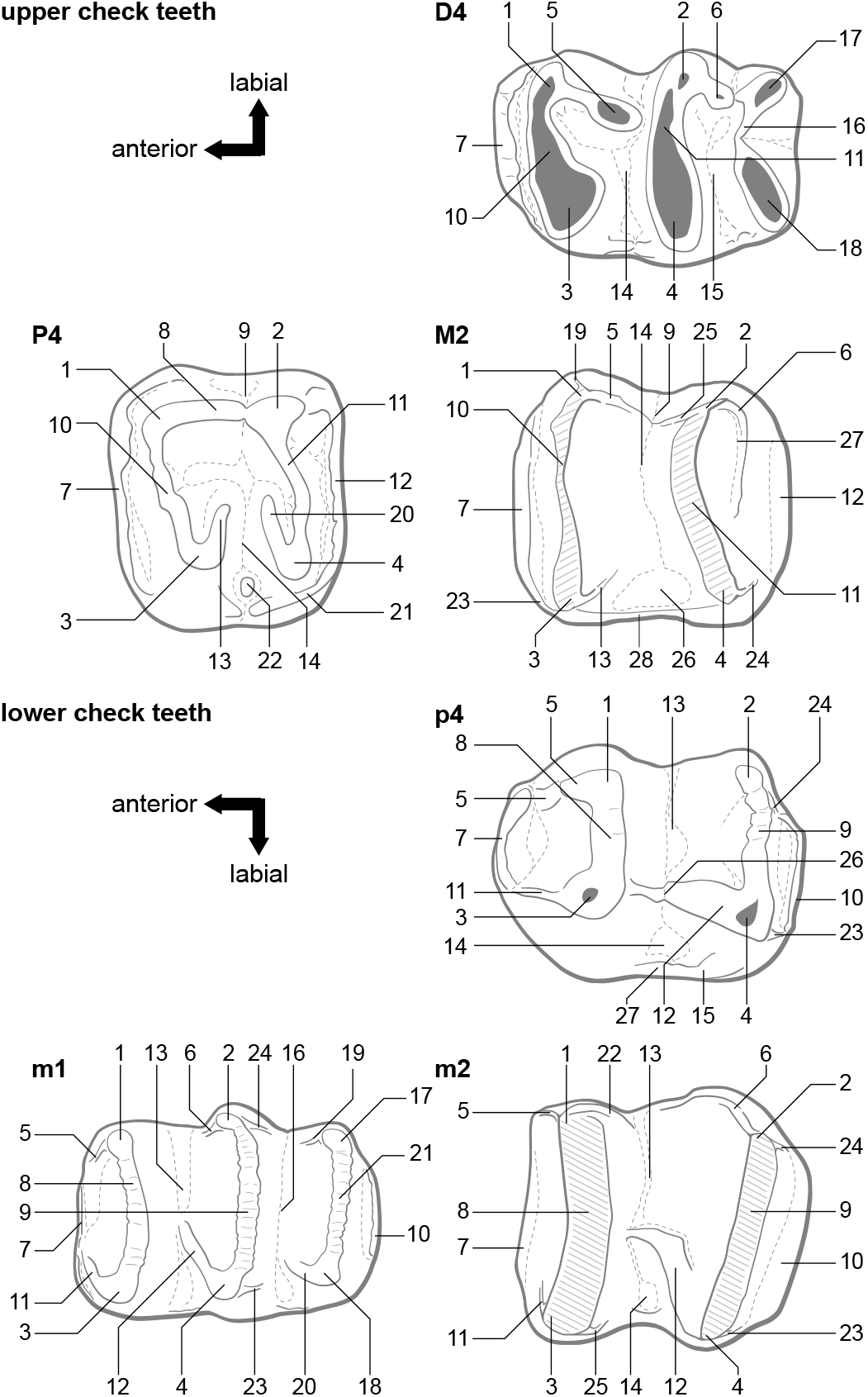
Dental terminology of upper and lower cheek teeth of Deinotheriidae in occlusal views (not to scale), mainly following Aiglstorfer et al. (2014) and Pickford and Pourabrishami (2013). **Upper cheek teeth**: 1, paracone; 2, metacone; 3, protocone; 4, hypocone; 5, postparacrista; 6, postmetacrista; 7, anterior cingulum; 8, ectoloph; 9, ectoflexus; 10, protoloph; 11, metaloph; 12, posterior cingulum; 13, postprotocrista; 14, median valley; 15, distal valley; 16, tritoloph; 17, labial tritoloph cone; 18, lingual tritoloph cone; 19, praeparacrista; 20, praehypocrista; 21, lingual cingulum; 22, entostyle (mesostyle of **Harris, 1973**); 23, praeprotocrista; 24, posthypocrista; 25, praemetacrista; 26, lingual medifossette; 27, convolute; 28, lingual cingulum. **Lower cheek teeth**: 1, metaconid; 2, entoconid; 3, protoconid; 4, hypoconid, 5, praemetacristid; 6, praeentocristid; 7, anterior cingulid; 8, metalophid; 9, hypolophid; 10, posterior cingulid; 11, praeprotocristid; 12, praehypocristid; 13, median valley; 14, labial medifossette; 15, labial cingulid; 16, distal valley; 17, lingual tritolophid conid; 18, labial tritolophid conid; 19, anterior cristid of the lingual tritolophid conid; 20, anterior cristid of the labial tritolophid conid; 21, tritolophid; 22, postmetacristid; 23, posthypocristid; 24, postentocristid; 25, postprotocristid; 26, labial notch; 27, labial cingulid.

### Systematics

The taxonomy of Deinotheriidae is still a debated issue as there is no consensus in the literature about the valid genera and species. Some authors point to very conservative morphological features of deinotheres and evolutionary changes essentially characterized by a gradual size increase through time, referring to *Deinotherium* as the only valid genus (e.g. **Gräf, 1957**; **Ginsburg and Chevrier, 2001**; **Pickford and Pourabrishami, 2013**). Others follow the two genera concept, *Prodeinotherium* and *Deinotherium*, as proposed by **Éhik (1930)**, based on dental, cranial and postcranial features (e.g. **Harris, 1973, 1978**; **Gasparik, 1993**; **Huttunen, 2002a**; **Huttunen and Göhlich, 2002**; **Duranthon et al., 2007**; **Vergiev and Markov, 2010**; **Aiglstorfer et al., 2014**; **Konidaris et al., 2017**; **Göhlich, 2020**). The recent data of European deinotheres support five different morphospecies or chronospecies (**Böhme et al., 2012**; **Pickford and Pourabrishami, 2013**), whereas previous investigations were favourable to four species (**Gasparik, 1993, 2001**; **Markov, 2008**; **Vergiev and Markov, 2010**) or even two species (**Huttunen, 2002a**). Resolving this issue is beyond the goal of this study. Hence, following the most recent publications on deinotheres such as **Aiglstorfer et al. (2014), Konidaris et al. (2017)** and **Göhlich (2020)**, this study refers to the two-genera taxonomic scheme and considers five European species to be valid: *Prodeinotherium cuvieri* (**Kaup, 1832**), *Prodeinotherium bavaricum* (**Meyer, 1831**), *Deinotherium levius* **Jourdan, 1861**, *Deinotherium giganteum* **Kaup, 1829** and *Deinotherium proavum* (**Eichwald, 1831**).

### Stratigraphy and fossil record

The stratigraphical framework used in this study is based on the global geological time scale for the Neogene (**Hilgen et al., 2012**), the European Mammal Neogene units (MN-Zones; **Mein, 1999**; **Steininger, 1999**), and the Swiss fauna references (**Engesser and Mödden, 1997**; **Berger, 2011**).

The data set of the fossil record of the European deinotheres is a compilation of the localities reported in **Maridet and Costeur (2010)**, The Paleobiology Database (extraction on the 09.08.2019 with the parameter family = Deinotheriidae; **PBDB, 2019**) and additional literature (**Supplementary information**). In order to highlight the palaeobiogeographic dynamics of distribution of deinotheres in Europe, localities are grouped by the biochronological intervals MN3-4, MN5-8, MN9-10 and MN11-13, and biogeographic events (Proboscidean Datum Event, Hipparion Datum Event) and major climate changes (Miocene Climatic Optimum, Mid-Miocene Cooling Event, Messinian Crisis) are taken into account. The biostratigraphical correlation of each locality was systematically checked in the literature and questionable data were removed from the data set.

## Abbreviations

APD: anteroposterior diameter
D: deciduous upper premolar
dex: right
H: height
i: lower incisors
L: length
m/M: lower and upper molars
MJSN: Jurassica Museum (formerly Musée jurassien des Sciences naturelles)
MN: Mammal Neogene
NMB: Naturhistorisches Museum Basel
NMBE: Naturhistorisches Museum Bern
p/P: lower and upper premolars
sin: left
TD: transverse diameter
W: width.

## Systematics

> Class MAMMALIA **Linnaeus, 1758**
>
> **Order PROBOSCIDEA Illiger, 1811**
>
> Family Deinotheriidae **Bonaparte, 1845**
>
> Genus *Prodeinotherium* **Éhik, 1930**

European species: *Prodeinotherium bavaricum* (**Meyer, 1831**), *P. cuvieri* (**Kaup, 1832**)

> ***Prodeinotherium bavaricum*** (**Meyer, 1831**)
>
> (**Figs 4–5; Tables 1–3**)
>
> Table 1. Dimensions [mm] of P4 and M2 of *Prodeinotherium bavaricum*. NMB-Mch.4, Montchaibeux, MN5-6; MJSN- VDL-001, Delémont valley, middle Miocene; NMB-D.G.5, Haute Garonne of Aurignac, middle Miocene; NMB-Fa.129, NMB-Fa.167, Pontlevoy-Thenay, MN5. 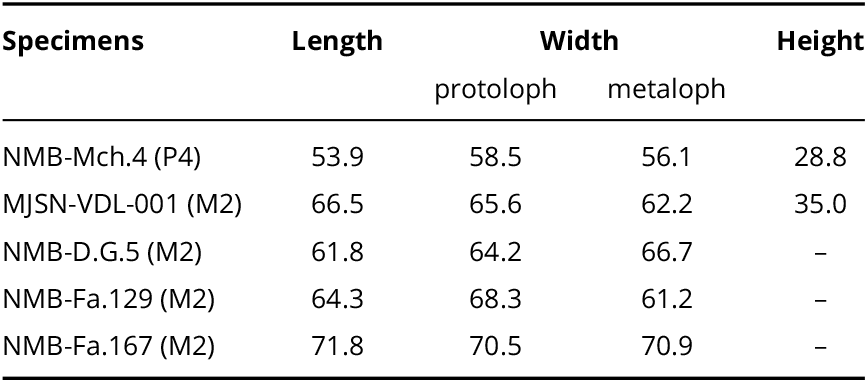
>
> Figure 4. *Prodeinotherium bavaricum* from the Delémont valley (Jura, Switzerland). **a**, P4 dex. (NMB-Mch.4, Montchaibeux locality) in labial (a1) and occlusal (a2) views; **b**, M2 dex. (MJSN-VDL-001, unknown locality) in occlusal (b1) and labial (b2) views; **c**, p4-m3 sin. (NMBE-5031977, Montchaibeux locality) in occlusal view. Scale bar = 5 cm. 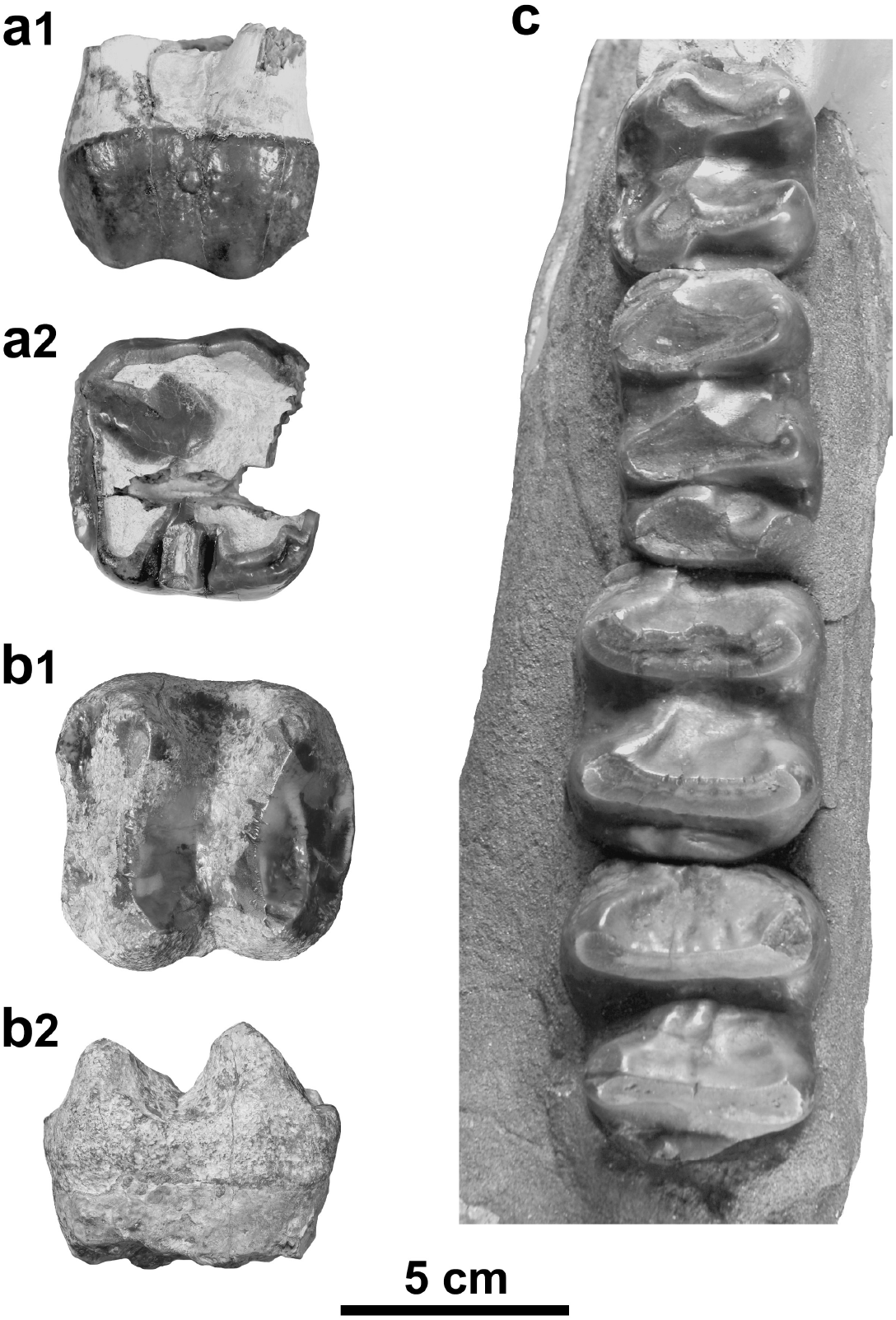
>
> Figure 5. *Prodeinotherium bavaricum* from Montchaibeux (Jura, Switzerland). **a**, Mandible (NMBE-5031977) in lateral view (a1), in anterior view (a2) and in dorsal view (a3). Scale bar = 20 cm. 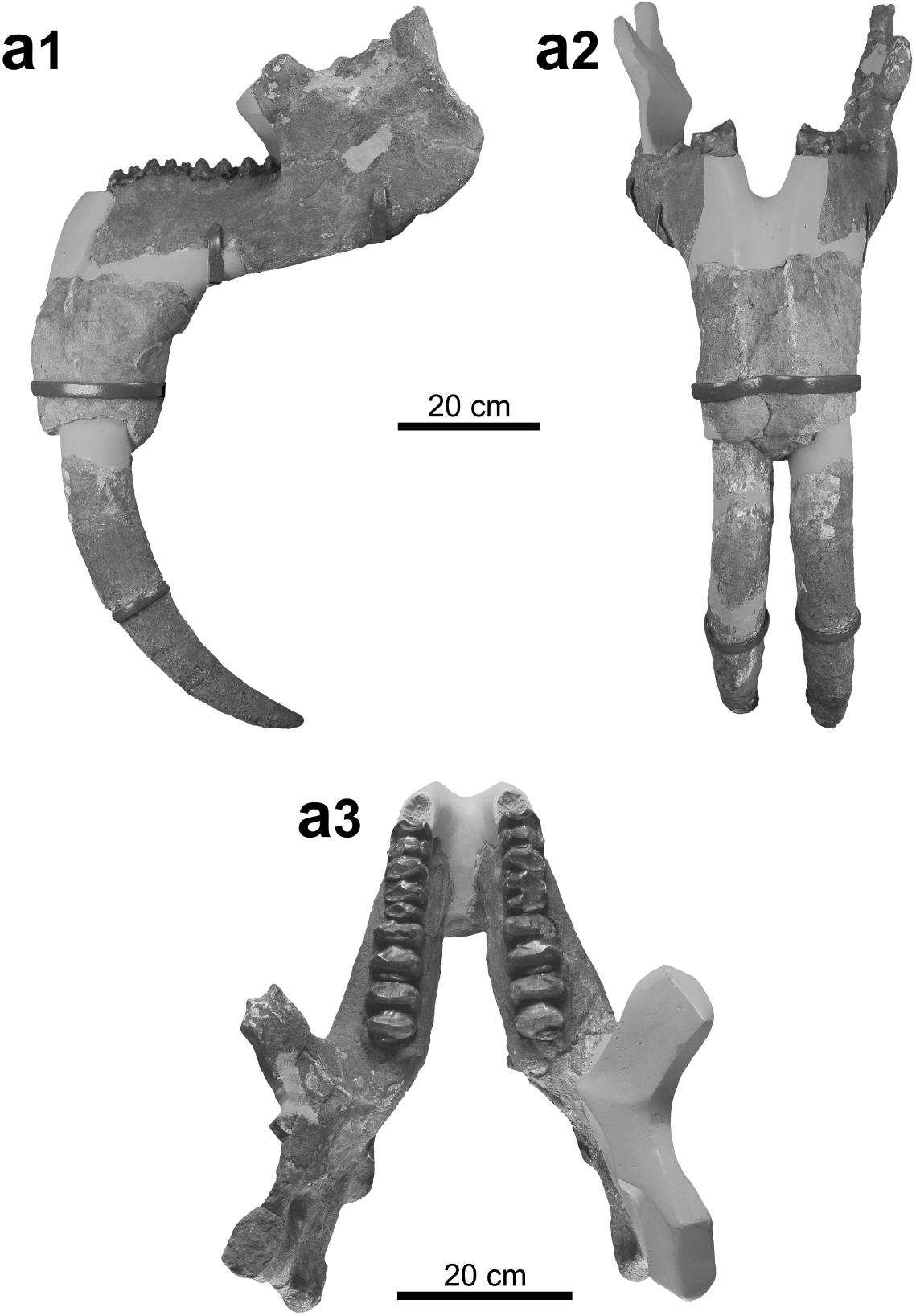

## Stratigraphical range

Late early and middle Miocene, MN4-6 (after **Gasparik, 2001**; **Huttunen and Göhlich, 2002**) or middle Miocene, MN5-6 (after **Pickford and Pourabrishami, 2013**; **Konidaris et al., 2017**; **Göhlich, 2020**).

## Material referred

M2 dex. (MJSN-VDL-001) from the Delémont valley (unknown locality); P4 dex. (NMB-Mch.4, copy MJSN- MTC-001) and mandible with i2 and p4-m3 (NMBE-5031977, copy MJSN-MTC-002) from Montchaibeux.

## Description

The P4 is damaged anteriorly and moderately worn. It is nearly quadratic in occlusal view, just slightly wider than long. The ectoloph is complete bearing an ectoflexus weakly developed, and distinct paracone fold, mesostyle (intermediate fold) and metacone fold, the former being the most developed. The protocone seems to extend labially, forming a complete protoloph reaching the paracone. The hypocone is labially elongated but does not form a complete metaloph connecting to the metacone, giving a sublophodont morphology to the tooth. The cingulum is posteriorly pronounced but anteriorly unobservable. The labial one is absent whereas the lingual one is strong but only present at the level of the protocone. The lingual opening of the median valley bears a well-developed entostyle. Three roots are present; the unique lingual one results from the fusion of two roots.

The M2 is bilophodont and subquadrate in occlusal view. The protoloph and metaloph are complete, both with almost the same width, anteriorly convex (with a more pronounced convexity on the metaloph) and have anterior wear facets. The four main cusps are distinct from the lophs. The postparcrista is wellmarked and slopes to the median valley that is opened on the lingual side. The postprotocrista is less developed and does not extend downward to the medial valley. The praehypocrista and the posthypocrista are not marked. The praemetacrista is well pronounced, slopes to the median valley and joins the postparacrista. Posteriorly, the convolute is well-developed. The anterior and posterior cingula are strong and continuous, although the anterior one is thinner in its middle part. The lingual cingulum is less pronounced but closes the lingual medifossette. The labial side of the tooth lacks any cingulum, but it is characterised by a deep ectoflexus.

The mandible NMBE-5031977, restored from five fragments, is incomplete. The ramus is low and slightly inclined forward, the mandibular angle and has an elevated position, the base of the corpus is straight, and the posterior margin of the symphysis is located below the front of the p4. The i2 are oriented downward and slightly curved backwards in their distal parts. The toothrows are almost complete from p4 to m3, the p4s being anteriorly incomplete and the p3 not preserved. The m1s are trilophodont and the other lower cheek teeth are bilophodont. The transverse lophids are subparallel, posteriorly convex for the anterior ones to straight for the posterior ones, and possess wear facets posteriorly oriented.

In occlusal view, the p4 is rectangular, longer than wide. The paracristid is not preserved, the metalophid is posteriorly convex and the hypolophid is almost straight. The ectolophid is poorly developed and descends anterolingually to reach the median valley. The labial cingulid is reduced to the posterior part of the tooth, the lingual one is lacking. The posterior cingulid is well developed, continuous and low but merging with a weak posthypocristid.

The rectangular m1 is trilophodont, with sub-parallel, roughly straight and of equally wide transverse lophids. The praeprotocristid, the praehypocristid and the anterior cristid of the labial tritolophid conid are all well pronounced, the latter two reaching the bottom of the respective front valleys. The anterior and posterior cingulids are poorly developed whereas the labial and lingual ones are lacking.

The m2 is sub-rectangular in occlusal view, slightly longer than wide, with equally wide transverse lophids. The metalophid is posteriorly slightly convex and the hypolophid is straight. The praeprotocristid and the praehypocristid are well developed and anterolingually oriented, the former reaching the bottom of the median valley. The anterior and posterior cingulids are continuous, the posterior one being stronger. The lingual and labial cingulids are lacking.

The m3 is morphogically similar to the m2. However, the hypolophid is slightly reduced in width compared to the metalophid and the posterior cingulid is more pronounced but strongly reduced in width, giving a longer and trapezoidal outline in occlusal view.

## Comparisons

The referred dental remains are typical of the Deinotheriidae family with mainly bilophodont jugal teeth associated to a sublophodont (well-developed ectoloph and incomplete metaloph) P4 and a trilophodont m1, as well as i2 oriented downwards and backwards (**Huttunen, 2002a**).

The specimens differ from *Deinotherium proavum* and *D. giganteum* by their considerably smaller dimensions (**Gräf, 1957**; **Tobien, 1988**; **Vergiev and Markov, 2010**; **Pickford and Pourabrishami, 2013**; **Aiglstorfer et al., 2014**; **T, ibuleac, 2018**). *Deinotherium levius* also presents larger dimensions (**Fig. 6**), but the differences are less significant as previously noticed in several studies (**Gräf, 1957**; **Tobien, 1988**; **Pickford and Pourabrishami, 2013**). However, the strong development of the convolute and the near absence of postprotocrista and posthypocrista on the M2 clearly exclude an attribution to *Deinotherium* (**Harris, 1973**; **Huttunen, 2002b**; **Poulakakis et al., 2005**; **Duranthon et al., 2007**; **Aiglstorfer et al., 2014**). Likewise the moderately developed curve of the i2 can be distinguished from the more pronounced one of *D. giganteum* and the subvertical one of *D. levius* (**Gräf, 1957**) and the position of the mandibular angle is more elevated than that of *Deinotherium* species (see **Gräf, 1957, fig. 12**; **Svistun, 1974, pl. 1**; **Huttunen and Göhlich, 2002, fig. 3**; **Vergiev and Markov, 2010, figs 3–4**; **Iliopoulos et al., 2014, fig. 1**).

**Figure 6.**
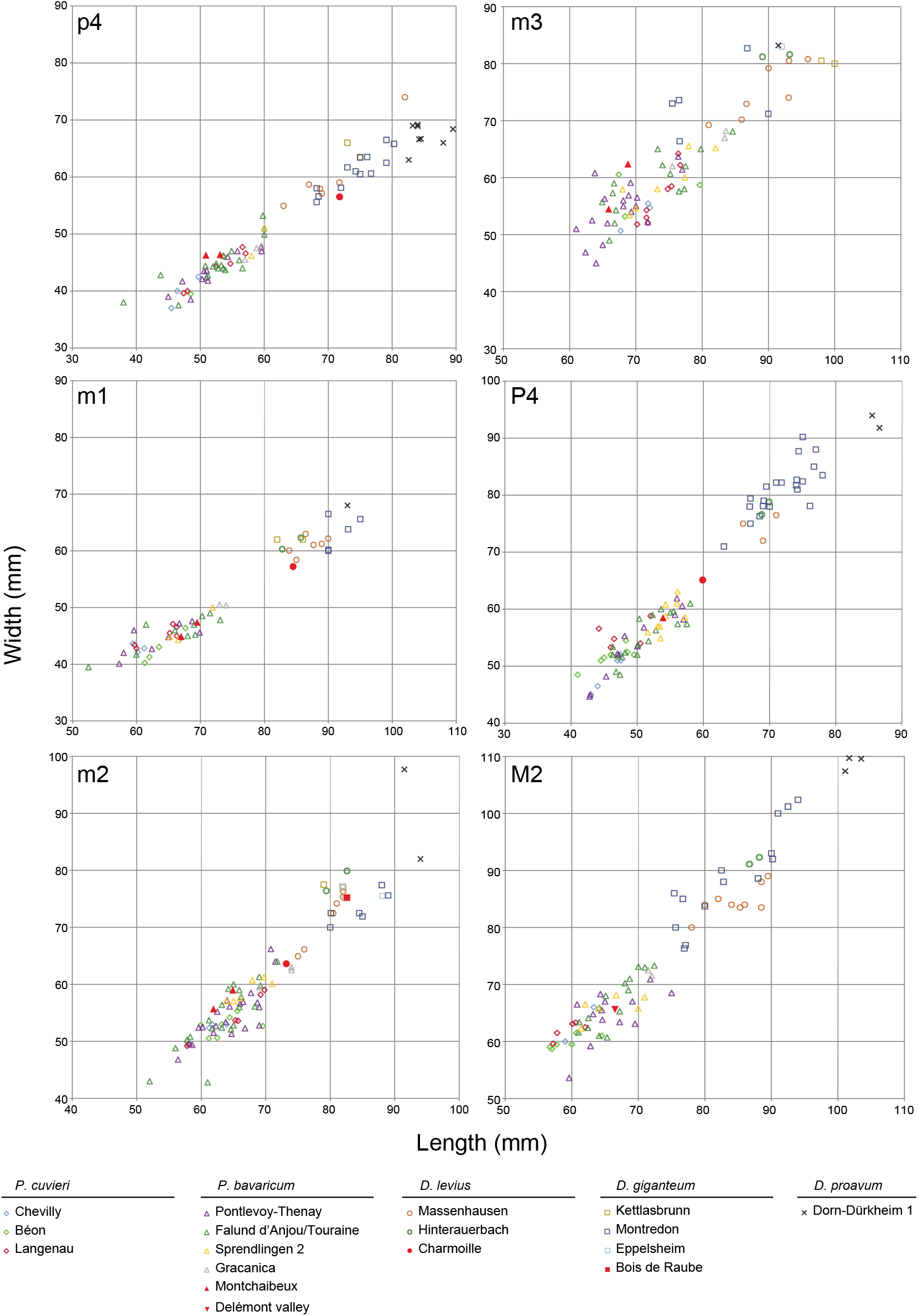
Squatter diagram of the teeth sizes (in mm) for p4, m1, m2, m3, P4 and M2 compared to deinotheres of other well-dated European localities. The measurements of Delémont valley (MN7/8), Montchaibeux (MN5-6), Charmoille (MN9) and Eppelsheim (MN9) come from this study; the measurements of Béon (MN4), Chevilly (MN4), Dorn-Dürkheim (MN11), Massenhausen (MN7/8) and Sprendlingen 2 (MN6) come from **Pickford and Pourabrishami (2013)**; the measurements of Falund d’Anjou/Touraine (MN5), Gracanica (MN5), Gratkorn (MN7/8), Hinterauerbach (MN7/8), Kettlasbrunn (MN9), Langenau (MN4), Montredon (MN10) and Pontlevoy-Thenay (MN5) come from **Göhlich (2020)**.

Although, in *Prodeinotherium*, the entostyle is usually lacking on P3-4 and the metaloph usually complete on P4, these particular characters, present on the referred P4 NMB-Mch.4, can be attributed to generic variability (e.g. **Harris, 1973**; **Ginsburg and Chevrier, 2001**; **Aiglstorfer et al., 2014**). Also by its dimensions, the almost absence of an ectolflexus and the quadratic outline in occlusal view, this specimen shows strong similarities with *P. bavaricum* (**Ginsburg and Chevrier, 2001**; **Duranthon et al., 2007**; **Pickford and Pourabrishami, 2013**). Based on the morphology of the P4 (nearly absence of ectolflexus and the quadratic outline), the M2 (developed convolute) and the lower cheek teeth (m1 with transverse lophids roughly straight and of equal width), as well as the modest curve of the i2 and the more elevated position of the mandibular angle, the specimens can be referred to the genus *Prodeinotherium* (**Gräf, 1957**; **Harris, 1973**; **Huttunen, 2002a**; **Huttunen and Göhlich, 2002**; **Duranthon et al., 2007**). Additionally, after **Ginsburg and Chevrier (2001)** and **Pickford and Pourabrishami (2013)**, the specimens cannot be referred to the species *P. cuvieri* due to their larger dimensions. Although the sizes of *P. cuvieri* and *P. bavaricum* show a lot of overlap, *P. bavaricum* remains on average larger (**Fig. 6**), as also noticed in previous studies (e.g. **Gräf, 1957**; **Kovachev and Nikolov, 2006**; **Huttunen and Göhlich, 2002**). Our specimens usually fit within the upper size-range of these measurements thus supporting a specific identification as *P. bavaricum* rather than *P. cuvieri*.

> Genus *Deinotherium* **Kaup, 1829**

European species: *Deinotherium giganteum* **Kaup, 1829**, *D. proavum* (**Eichwald, 1831**), *D. levius* **Jourdan, 1861**.

> ***Deinotherium levius* Jourdan, 1861**
>
> (**Fig. 7; Table 4**)
>
> Table 2. Dimensions [mm] of the mandible of *Prodeinotherium bavaricum* (NMBE-5031977) from Montchaibeux (Jura, Switzerland, MN5-6). 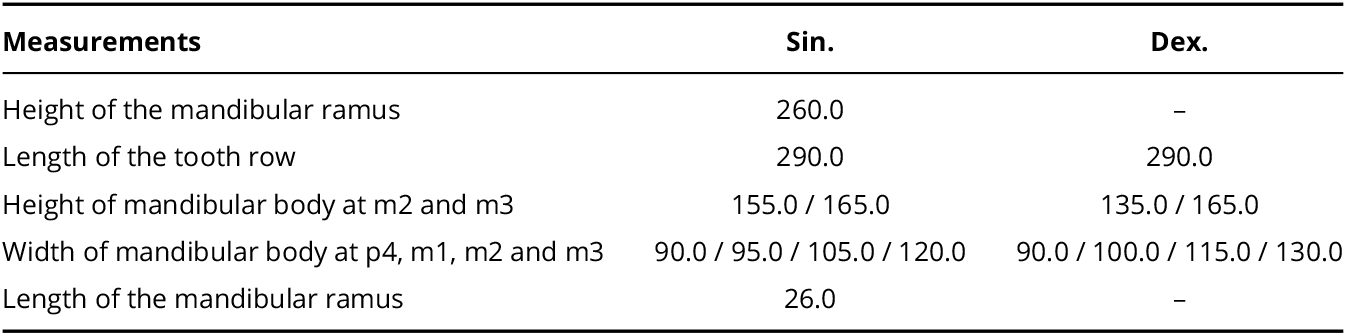
>
> Table 3. Dimensions [mm] of the teeth of the mandible of *Prodeinotherium bavaricum* (NMBE-5031977) from Montchaibeux (Jura, Switzerland, MN5-6). 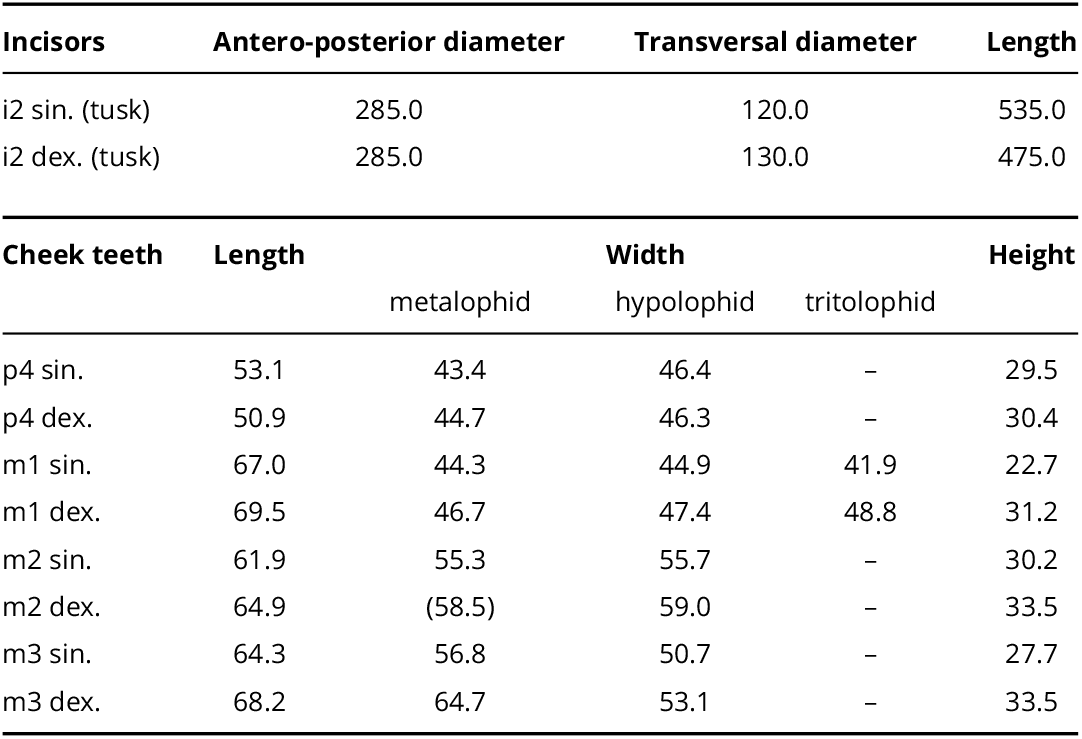
>
> Table 4. Dimensions [mm] of the teeth of *Deinotherium levius* from Charmoille (Jura, Switzerland, MN9). 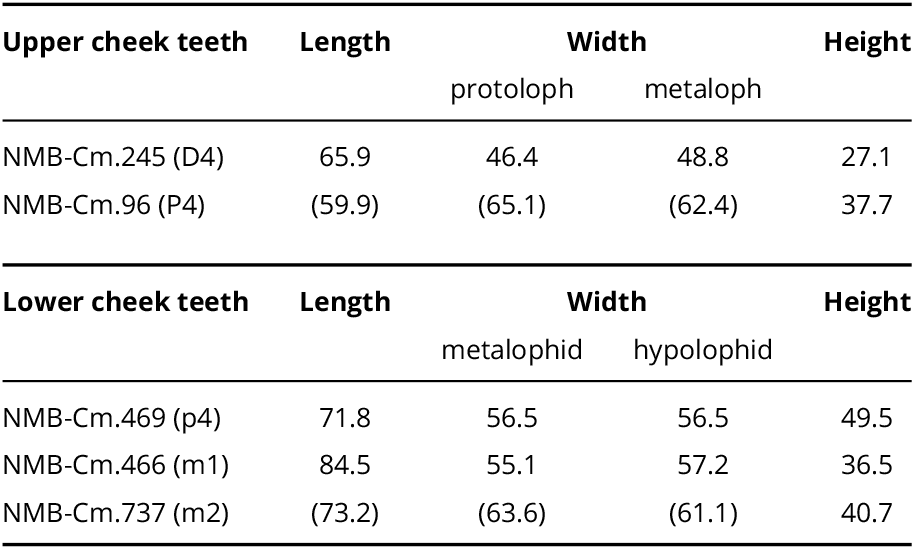
>
> Figure 7. *Deinotherium levius* from Charmoille (Jura, Switzerland). **a**, D4 dex. (copy MJSN-CH-060 of NMB- Cm.245,) in labial (a1) and occlusal (a2) views; **b**, P4 dex. (copy MJSN-CH-062 of NMB-Cm-96,) in labial (b1) and occlusal (b2) views; **c**, p4 dex. (copy MJSN-CH-058 of NMB-Cm.469,) in occlusal (c1) and labial (c2) views; **d**, m1 dex. (copy MJSN-CH-059 of NMB-Cm.466,) in occlusal (d1) and labial (d2) views; **e**, m2 dex. (copy MJSN-CH-061 of NMB- Cm.737,) in occlusal (e1) and labial (e2) views. For better illustration quality, white copies were photographed. Scale bar = 5 cm. 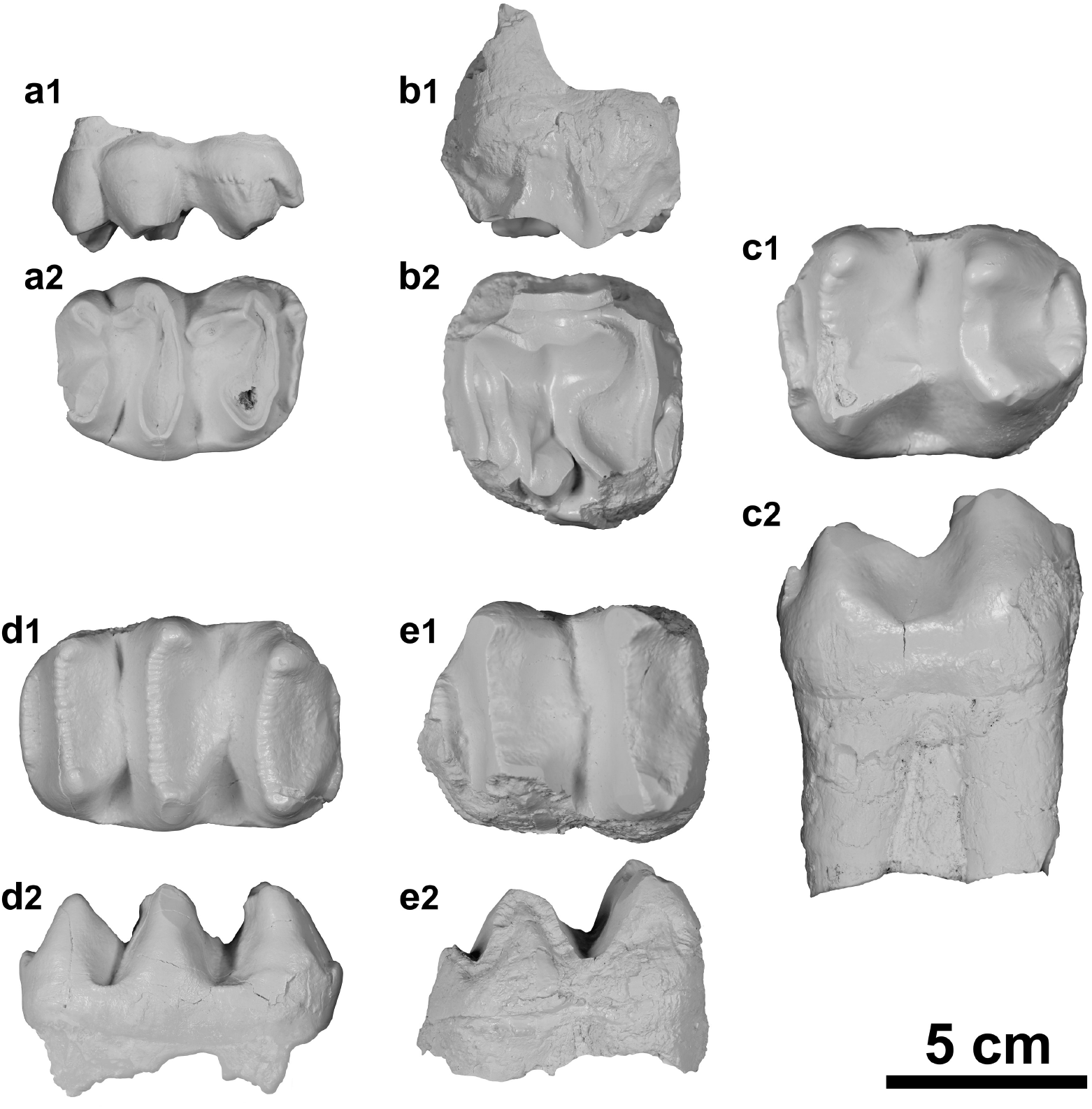

## Stratigraphical range

Late middle to early late Miocene MN7/8-9 (**Göhlich and Huttunen, 2009**; **Aiglstorfer et al., 2014**; **Konidaris et al., 2017**; **Konidaris and Koufos, 2019**; this study).

## Material referred

Distal fragment of a right incisor (NMB-Cm.478), D4 dex. (NMB-Cm.245, copy MJSN-CH-060), P4 dex. (NMB-Cm-96, copy MJSN-CH-062), p4 dex. (NMB-Cm.469, copy MJSN-CH-058), m1 dex. (NMB-Cm.466, copy MJSN-CH-059) and m2 dex. (NMB-Cm.737, copy MJSN-CH-061) from Charmoille in Ajoie.

## Description

The fragmented incisor NMB-Cm.478 is roughly oval in transverse section, with a longest axis in anteroposterior direction, the diameter diminishing distally and a flattened medial side. The distal curvature, caudally and laterally, is weakly developed. The specimen shows wear facets on the distal side and at the tip.

The D4 is trilophodont and elongated. The protoloph is anteriorly convex and the metaloph is nearly straight. The tritoloph is anteriorly strongly convex and incomplete; the lingual and labial cones are separated by a notch. The postparacrista and the postmetacrista are well-developed, extending posterolingually downward and reaching the rear loph. The anterior and posterior cingula are present, the anterior one being strongly pronounced and connected to the paracone by a faint crista. The transverse valleys are lingually faintly closed by a reduced lingual cingulum. The labial cingulum is almost completely lacking, only faint labial rugosities are observable at the level of the paracone.

The P4 is moderately worn, incomplete (enamel only partly preserved around the outline of the crown), slightly wider than long and trapezoidal in occlusal view. The ectoflexus is very smooth and the mesostyle barely distinct. The protoloph is complete, reaching the paracone, whereas the metaloph is in contact with the metacone but not fused with it. The hypocone extends anterolabially downward by a praehypocrista. The cingulum is absent labially, is anteriorly and posteriorly strong and continuous, and is labially reduced to the opening of median valley. The latter bears a strong entostyle in contact with the hypocone but separated from the protocone. The two lingual roots are isolated and the two lingual ones are in contact, just separated by a vertical groove.

The p4 is almost bilophodont with the occlusal outline longer than wide. An ectolophid extends anterolingually downward from the hypoconulid, reaching the base of the metalophid. The metalophid is anteriorly concave, the hypolophid is roughly straight. The paracristid extends anteriolingually downward from the paraconid and connects a very strong anterior cingulid. The praemetacristid extends anteriorly downward, almost closing an anterior valley-like groove. The posterior cingulid is well developed and connected to the hypoconulid by a very faint posthypocristid. The lingual cingulid is lacking and the labial one is reduced to the base of the labial notch, closing a labial medifossette.

The m1 is trilophodont and elongated. Each conid has a slightly pronounced anterior cristid. The praehypocristid is the most developed. It extends anterolingually downward, reaching the anterior valley and reaching the metalophid. The anterior cingulid is poorly developed whereas the posterior one is more developed. The transverse valleys are open on both sides, although reduced labial cingulids are present at the extremities of these valleys.

The m2 is bilophodont and nearly rectangular (slightly longer than wide). The anterior cingulid is unobservable whereas the posterior one is low and strong but narrower than the hypolophid. The median valley is opened on both sides, without labial and lingual cingulids. Each conid has a slightly developed and anteriorly extending cristid, except the praehypocristid which extends anterolingually and reaches the bottom of the median valley.

## Comparisons

The specimens from Charmoille show the typical features of Deinotheriidae: lower tusks oriented downward, P4 bearing an ectoloph, trilophodont D4 and m1, and a bilophodont pattern for the remainder of the cheek teeth (**Huttunen, 2002a**). They differ from *Prodeinotherium* by being larger, by a trapezoidal outline and a more distinct ectoflexus in P4, as well as a narrower tritolophid compared to other lophids in m1 (**Gräf, 1957**; **Ginsburg and Chevrier, 2001**; **Duranthon et al., 2007**). Among the *Deinotherium* species, they display more affinities with *D. levius* by the size (**Figs 6, 8**; i.e., larger than *P. cuvieri* and *P. bavaricum* and slightly smaller than *D. giganteum*; **Pickford and Pourabrishami, 2013**), by a subcomplete metaloph without a notch separating it from the metacone and the presence of a strong entostyle on P4, by a protolophid and metalophid of equal lengths in p4 (rectangular outline vs trapezoidal outline in *D. giganteum*), and by a short posterior cingulid on m2 (**Gräf, 1957**; **Duranthon et al., 2007**). This attribution is also supported by the i2 NMB-Cm478 that displays a sub-straight tusk tip, characteristic of *D. levius* according to **Gräf (1957)**.

**Figure 8.**
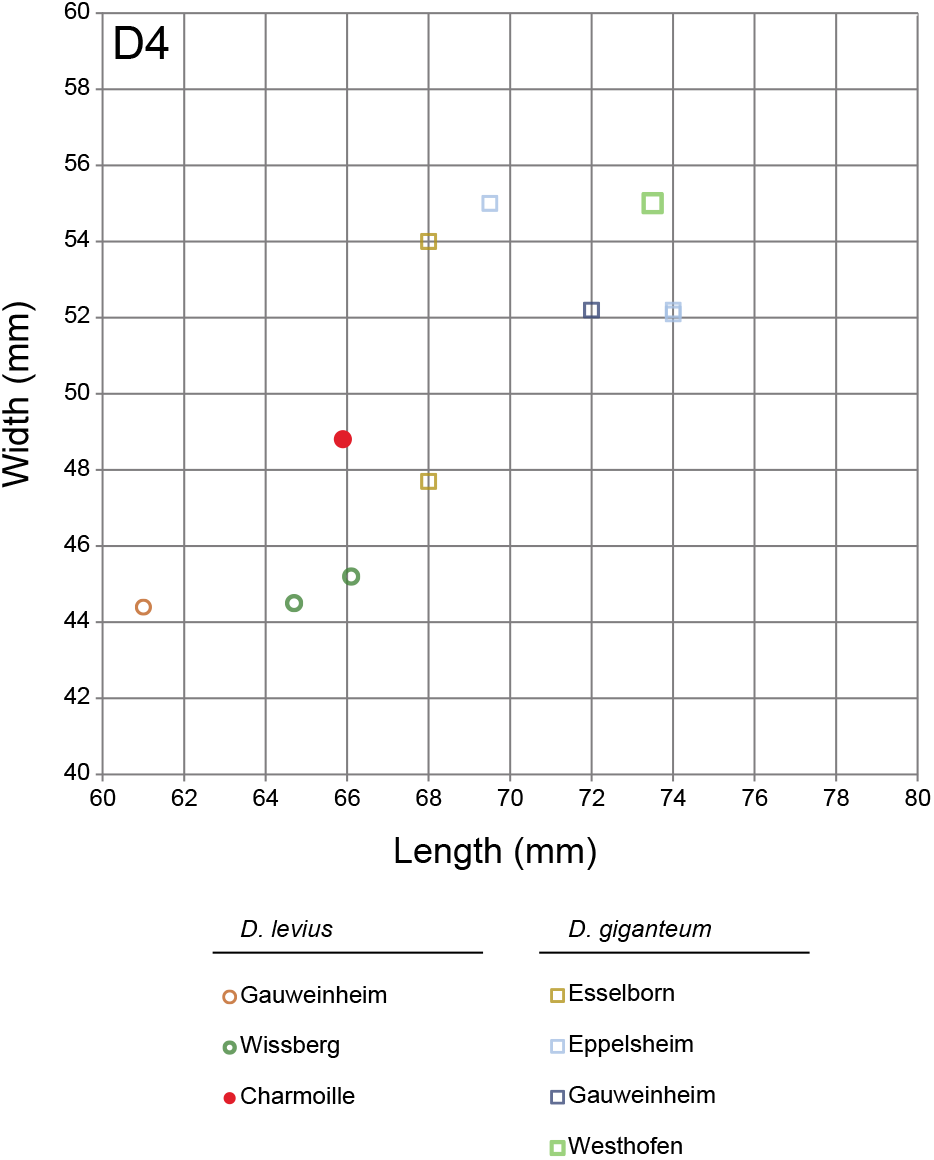
Squatter diagram of the teeth sizes (in mm) for D4 compared to deinotheres of other well-dated European localities. The measurements of Charmoille (MN9) come from this study; the measurements of Gauweinheim (MN7/8), Esselborn (MN9), Westhofen (MN9) and Wissberg (MN7/8) come from **Pickford and Pourabrishami (2013)**.

> ***Deinotherium giganteum* Kaup, 1829**
>
> (**Fig. 9; Table 5**)
>
> Figure 9. *Deinotherium giganteum* from Bois de Raube in the Delémont valley (Jura, Switzerland). **a**, m2 sin. (copy MJSN-BRA-001) in lingual (a1), occlusal (a2) and labial (a3) views. For better illustration quality, a white copy was photographed. Scale bar = 5 cm. 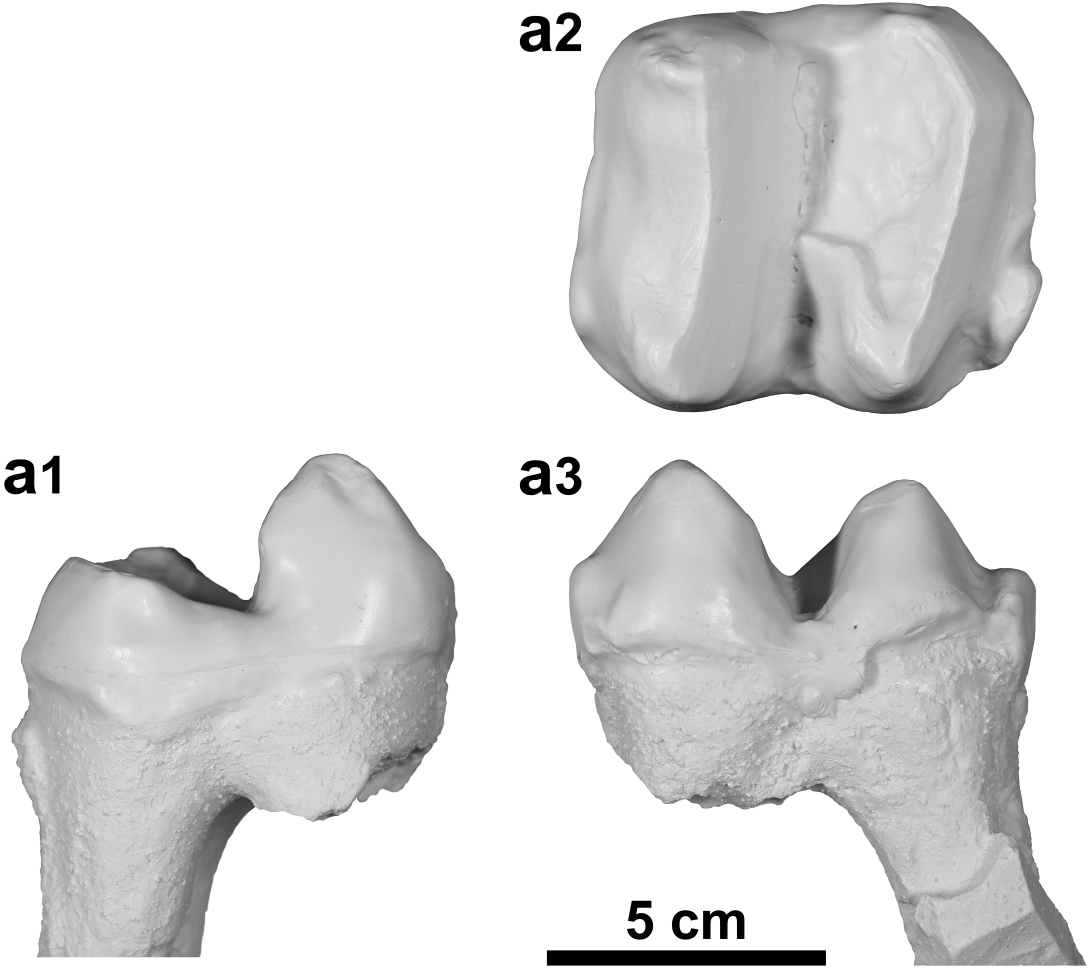
>
> Table 5. Dimensions [mm] of m2 of *Deinotherium giganteum* (copy MJSN-BRA-001, Bois de Raube, Jura, Switzerland, MN6-7/8; NMB-Ep.16, NMB-Ep.135, Eppelsheim, Germany, MN9). 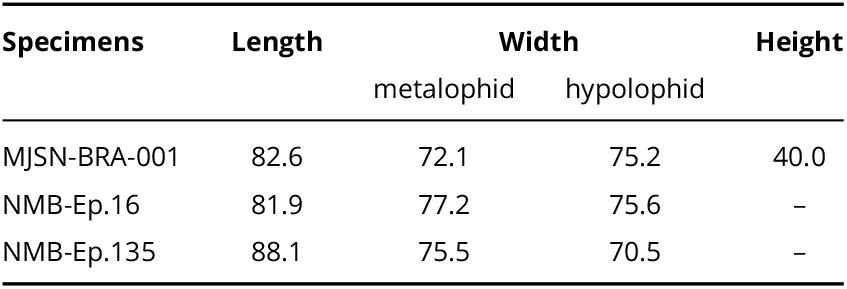

## Stratigraphical range

Late middle to early late Miocene MN7/8-10 (**Konidaris et al., 2017**; this study).

## Material referred

Complete m2 sin. (copy MJSN-BRA-001; original in the Jean-Baptiste Greppin collection at the Strasbourg University) from the Bois de Raube locality in the Delémont valley.

## Description

The referred m2 is bilophodont and slightly longer than wide in occlusal view. The four main cuspids are distinct. The transverse lophids are complete, separated by a labially deeper median valley, and have posteriorly wear facets. The hypolophid is sublinear and slightly wider than the metalophid. The metalophid is anteriorly weakly concave. The protoconid and the metaconid are quite sharp and equally height. The prae- and postprotocristid are hardly distinct, the prae- and postmetacristid are more prominent but blunt. The entoconid is very smooth, difficult to distinguish and lower than the metaconid. The praeentocristid is quite well marked, very rounded, and descends almost to the level of the median valley. The postentocristid is barely visible. The hypoconid, quite salient at the top, is slightly more modest than the protoconid. The praehypocristid, really robust and smooth, forms a thick enamel bulge that descends transversally to the median valley level and almost reaches the middle of the tooth. The posthypocristid is very weak, almost indistinct. There is no particular ornamentation on the tooth. However, the presence of a strong posterior cingulid, incomplete on the labial side, of a weak anterior cingulid, slightly more pronounced labially, and of a labial medifossette barely delimited by a modest enamel bridge are noticeable.

## Comparisons

The m2 displays a bilophodont pattern with a well-developed posterior cingulid which are typical of the Deinotheriidae family (**Huttunen, 2002a**). This m2 can be differentiated from m2s of *Prodeinotherium* by their sizes (**Fig. 6**) that are on average up to more than 30% larger than those of *P. cuvieri* and about 20% larger than those of *P. bavaricum* (e.g. **Gräf, 1957**; **Ginsburg and Chevrier, 2001**; **Huttunen and Göhlich, 2002**; **Pickford and Pourabrishami, 2013**). In addition, the praehypocristid is remarkably more developed than in *P. bavaricum* (as is the posterior cingulid too), then the tooth can undoubtedly be referred to the genus *Deinotherium* (e.g. **Huttunen, 2002a, b**; **Huttunen and Göhlich, 2002**; **Duranthon et al., 2007**; **T, ibuleac, 2018**).

A specific identification within the genus *Deinotherium* remains very difficult based on morphological characters whereas size increase seems to be most obvious change interpreted as an evolutionary trend through time (e.g. **Gräf, 1957**; **Ginsburg and Chevrier, 2001**; **Duranthon et al., 2007**; **Pickford and Pourabrishami, 2013**). However, **Pickford and Pourabrishami (2013)** suggest specific attributions by highlighting, contrary to **Gräf (1957)**, discontinuous size ranges from one species to another. Based on these observations, m2s of *D. proavum* are always larger than 90 mm and can exceed 100 mm, which unambiguously excludes our specimen from Bois de Raube whose length is 82.6 mm (**Table 5**). The m2 of Bois de Raube (MJSN-BRA-001) falls within the length-range of *D. giganteum* between 70.0 and 89.0 mm but also corresponds to the largest size of *D. levius* (**Fig. 6**). However, *D. levius* remains on average smaller than *D. giganteum* in which size-range the m2 from Bois de Raube fits better, also the degree of development of the posterior cingulid shows a very close similarity to m2 of *D. giganteum* from Eppelsheim (NMB-Ep.16, NMB-Ep.135) and from Romania (**T, ibuleac, 2018**). For these reasons, we refer this isolated tooth to *D. giganteum*.

## Discussion

### Fossil record of Deinotheriidae in the Jura

The age of the deinotheres discovered in the Swiss Jura Mountains is based on the regional litho- and biostratigraphy established by **Kälin (1993, 1997)** and **Prieto et al. (2018)** and fits the biostratigraphic range of the species at the European scale. The records correlate to MN5-6(−7) for *P. bavaricum* in Montchaibeux, to MN7/8 for *D. giganteum* in Bois de Raube and to MN9 for *D. levius* in Charmoille. The latter record indicates the first report of *D. levius* in Switzerland and matches the latest occurrences of this species in Europe (**Göhlich and Huttunen, 2009**; **Aiglstorfer et al., 2014**; **Konidaris and Koufos, 2019**), whereas the record of *D. giganteum* in Bois Raube could be among the youngest record of this taxa in Europe (**Fig. 10**).

**Figure 10.**
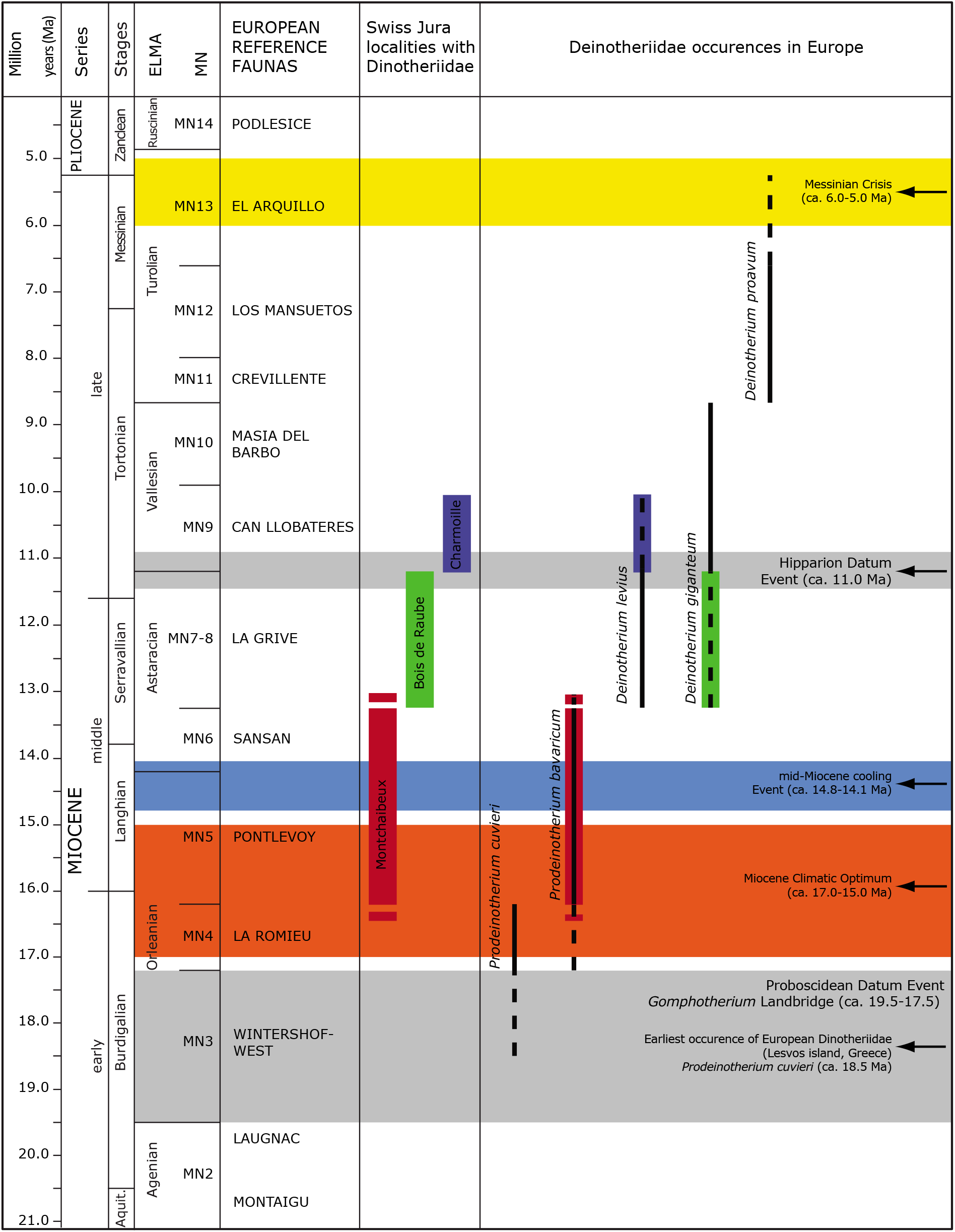
Stratigraphic extent of the five European species of Deinotheriidae (*P. cuvieri, P. bavaricum, D. levius, D. giganteum* and *D. proavum*). The dashed lines represent enlarged occurrences for each species, supported by the fossil record of the **Supplementary information**. The correlations with the European fauna of reference are according to **Berger (2011)** and the ones with the regional lithostratigraphy according to **Kälin (1993, 1997)** and **Prieto et al. (2018)**.

### Biogeographic distribution of European Deinotheriidae

The deinotheres known since the late Oligocene in Africa arrived later in Eurasia, following the mid-Burdigalian Proboscidean Datum Event (ca. 19.5-17.5 Ma). This event is related to the terrestrial corridor, called the *Gomphotherium* Landbridge, allowing a faunal exchange between Eurasia and the Arabian Plate of which the proboscideans were the palaeontological index fossils (**Tassy, 1990**; **Göhlich, 1999**; **Rögl, 1999a, b**; **Koufos et al., 2003**). Although the first, short-lasting migration corridors evolved already during the Aquitanian or perhaps earlier in Asia (e.g. **Tassy, 1990**; **Antoine et al., 2003**), the main wave of migration of the *Gomphotherium* Landbridge started during the mid-Burdigalian in Europe, with the arrivals of the earliest gomphotheres, deinotheres and mammutids at the end of MN3 (**Tassy, 1990**; **Koufos et al., 2003**). Among the early occurrences of European deinotheres in MN3-4, *Prodeinotherium cuvieri* is better represented in the west of Europe (France and Spain; **Fig. 11**) except for the earliest occurrence in Lesvos Island (MN3; identified as *P. bavaricum* in **Koufos et al., 2003**, but corrected as *P. cuvieri* following the concept of the five valid European species as in **Aiglstorfer et al., 2014, Konidaris et al., 2017** and **Göhlich, 2020**) which is likely a record of the immigration itself. *Prodeinotherium bavaricum* could already be recorded as early as MN4, only in Hungary (identified as *P. hungaricum* by **Éhik, 1930** and **Gasparik, 1993, 2001**), then display a more even distribution over Europe since MN5 (**Fig. 11**). This period mainly corresponds to the Miocene Climatic Optimum (ca. 17.0-15.0 Ma) when a tropical forest covered most of Europe with an average annual temperature that could reach 20-22°C and a more marked seasonality (nearly six months of drought; **Böhme, 2003**).

**Figure 11.**
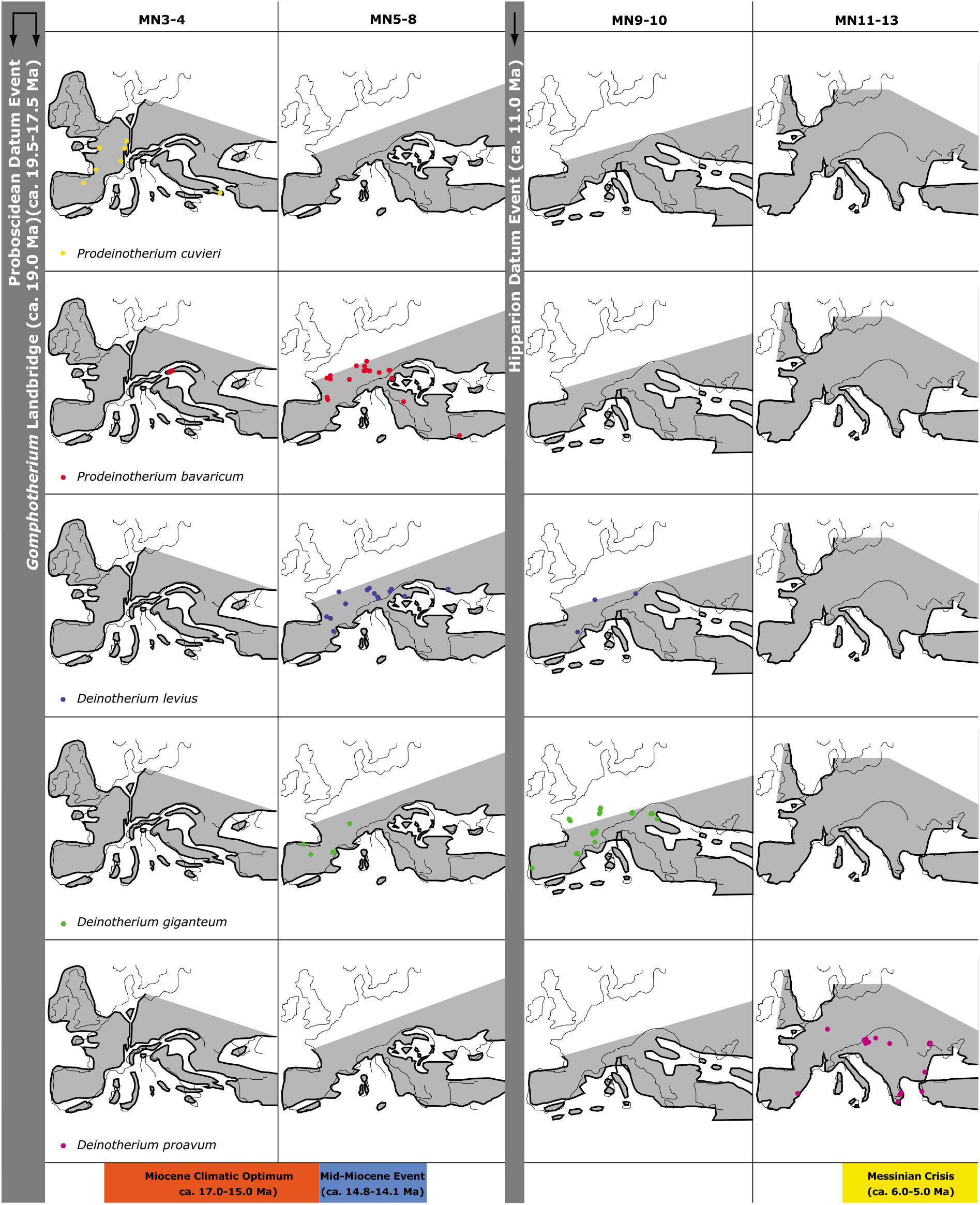
Palaeobiogeographic distribution of the five Deinotheriidae species in Europe (*Prodeinotherium cuvieri, P. bavaricum, Deinotherium levius, D. giganteum, D. proavum*). See localities in the **Supplementary information**.

After the Hipparion Datum Event (ca. 11.0 Ma), i.e. arrival in Europe of the little tridactyl horse from northern America (*Hippotherium primigenium*) throughout the Holarctic regions (**MacFadden, 1992**), the deinotheres are essentially dominated by *D. giganteum* during the Vallesian (e.g. **Göhlich, 2020**), although some rare occurrences *D. levius* are still reported (e.g. **Göhlich and Huttunen, 2009**; this study), and the huge *D. proavum* during the Turolian (e.g. **Konidaris et al., 2017**). The occurrence of *Deinotherium* is also confirmed at the latter time into the Middle East with *D. proavum* (most likely misidentified in *D. giganteum* by **Mirzaie Ataabadi et al., 2011**) and in India with *D. indicum* (**Sankhyan and Sharma, 2014**; **Singh et al., 2020**).

Finally, during the MN13 biozone, corresponding to the Messinian Crisis (ca. 6.0-5.0 Ma) and the extension of the open forests in the temperate latitudes of Eurasia (**Vislobokova and Sotnikova, 2001**; **Rouchy et al., 2006**), only the last representatives of *D. proavum* subsist in Eastern Europe (e.g. **Gasparik, 2001**).

### Morphological evolution and ecology of the Deinotheriidae

Teeth of Deinotheriidae show a remarkable increase of their dimensions throughout their evolution (**Pickford and Pourabrishami, 2013**) which reflects an evolution toward larger size for the whole family (**Aiglstorfer et al., 2014**; **Codrea and Margin, 2009**). According to **Agustí and Antón (2002)**, *Prodeinotherium* was 2 metres tall at the shoulder, while *Deinotherium* might have reached 4 metres. Some species of Deinotheriidae presented body mass far greater than those of extant elephants. For comparison, the greatest recorded weight of an African elephant is 6.64 tons (**Larramendi, 2016**), whereas the average ranges between 4 and 5 tons. The most ancestral deinotheres, *Chilgatherium harrisi*, weighed already 1.5 tons (**Sanders et al., 2004**), *Prodeinotherium bavaricum* and *P. hobleyi* weighed nearly 4 tons, *Deinotherium bozasi* about 9 tons, *D. levius* about 10 tons, while *D. giganteum* and *D. proavum* greatly exceeded 10 tons (**Larramendi, 2016**). All the Deinotheriidae representatives are therefore mega herbivores, i.e. mammals that feed on plants and reach a body mass of at least a ton or more at an adult age (**Owen-Smith, 1988**). Throughout the evolution of terrestrial mammals, a maximal limit of body mass of the mega herbivores could be of approximately 17 tons, estimated weight for *Paraceratherium transouralicum* (Rhinocerotoidea of the lower Oligocene in Eurasia) and some specimens of *Deinotherium* from the late Miocene of Eurasia and Africa (**Smith et al., 2010**). Nowadays, mega herbivores include elephants, most of rhinoceros, hippopotamus and giraffes, however none of these mammals reach 10 tons (**Owen-Smith, 1988**).

The body size and mass of mammals is linked to a large number of physiological and ecological traits (e.g., **Blueweiss et al., 1978**; **Brown et al., 2004**). The lifestyle, the living environment and the spatial distribution of the species are parameters particularly linked to the size (for a synthesis see **McNab, 1990** and **Eisenberg, 1990**). Having a large body size and mass brings consequently non-negligible advantages for the survival of a population, such as a lower mortality rate, a more stable population dynamic and a better resistance to sickness and limiting environment factors (**Erb et al., 2001**; **Langer, 2003**). Among large mammals, the mega herbivores are more immunised against predation thanks to their huge size and mass, providing also a protection to the youngest because of their generally gregarious behaviour (**Hummel and Clauss, 2008**). This advantage might have been particularly important during the Miocene that also sees a significant size augmentation of some predators (e.g., *Hyainailouros sulzeri, Amphicyon giganteus, Machairodus giganteus*; **Agustí and Antón, 2002**). Due to the opening of environments during the Neogene (e.g. **Suc et al., 1999**; **Favre et al., 2007**; **Costeur et al., 2007**; **Costeur and Legendre, 2008**), the folivore herbivores, such as the deinotheres, also had to browse over extended ranges from a wooded patch to another to find food. Large mammals have greater potential for long range dispersal and hence larger geographical distribution (e.g. **Brown, 1995**; **Gaston, 2003**), the displacements demanding less energy per distance unit for large animals (**Owen-Smith, 1988**). More important size and mass were therefore favourable in the environmental context of the Miocene in Europe. Lastly, the appearance of the first really large European species of *Deinotherium* (*D. levius, D. giganteum*) occurred in the middle Miocene, corresponding to the global fall of temperatures (Mid-Miocene Cooling Event, ca. 14.8- 14.1 Ma; **Flower and Kennett, 1994**). According to the Bergmann’s Law (**Bergmann, 1847**; **Blackburn and Hawkins, 2004**), although this rule suffers from numerous exceptions (**Meiri and Dayan, 2003**), a large body mass also allows a limitation of heat loss and presents a significant advantage in a colder climate. All these advantages linked to large size and mass could have supported the natural selection of larger deinotheres and in turn could explain the regular augmentation of size of this family during the Neogene.

The structure of the cheek teeth of deinotheres is specifically bilophodont and closer to those of tapirs than the multilophodont (lamellae) structure of extant elephants. Tapirs are essentially folivores and spend up to 90% of their active time to feed on fruits, leaves, barks and flowers (**Huttunen, 2002a**; **Naranjo, 2009**; **Sanders, 2020**). Likewise, deinotheres seem to be specialised in a regime consisting of dicotyledonous foliage and are generally linked to closed environmental patches (**Calandra et al., 2008**; **Čkonjevic and Radovic, 2012**; **Aiglstorfer et al., 2014**). Additionally, the gradual size increase observed in deinotheres through time (e.g. **Pickford and Pourabrishami, 2013**) seems to be associated to general evolution of environments in Europe: from rather closed forest environments of the early and middle Miocene and to rather open forest environments of the late Miocene (**Eronen and Rössner, 2007**).

In the more derived representatives of *Deinotherium*, the occiput is slightly inclined backwards and the occipital condyles elevated, characterizing a higher head posture (e.g. **Harris, 1973**). The appendicular skeleton also presents a modification of the graviportal structure initially known in *Prodeinotherium* leading to a more agile anatomic type with notably a greater amplitude of movements for the anterior limbs (scapular spine without acromion and metacromion, functional tetradactyly with a reduction of the first metacarpal and first metatarsal; **Huttunen, 2002a**). Therefore the association of the body size and mass and the anatomic evolution of Deinotheriidae suggest an ecological evolution at a family level, favouring the more mobile and larger species, adapted to more open and scattered forest landscapes. Such an evolutionary history could explain the progressive displacement of Deinotheriidae during the Miocene to Eastern Europe, where a drier climate (**Eronen et al., 2010**; **Bruch et al., 2011**) had probably favoured this type of environment.

## Conclusion

During the MN4-6 interval, only the small-sized deinotheres (*Prodeinotherium* species) are present, mostly in Western Europe. The occurrence of large sizes is recorded since MN7 with the genus *Deinotherium*. This genus shows a gradual size increase through time (MN7/8 to MN13) from *D. levius* and *D. giganteum* to *D. proavum*. The last deinotheres becomes gradually mostly restricted to Central and Eastern Europe, which seems to serve as a refuge area where only the huge eventually *D. proavum* eventually remains by the end of the Miocene.

Two factors seems to explain this palaeobiogeographic dynamic, the climatic evolution during the Miocene leading to a differentiation between the environments of Western and of Eastern Europe, and the ecological evolution of the deinothere. Since the beginning of the Miocene, Europe underwent numerous climatic changes that divided the continent in two really distinct environments from the late Miocene onward. Indeed, Western Europe environments were dominated by still closed and semi-humid tropical forests whereas Eastern Europe had more open and drier forest landscapes due to a more continental climate (**Vislobokova and Sotnikova, 2001**). Deinotheres being folivores were clearly linked to forest environments and needed large quantities of foliage throughout the year to sustain the amount of energy that their huge body mass required. The combination of their specialised diet and morphologic evolution (higher head posture, increased size and improved agility) reflects a remarkable adaptive and ecologic evolution of the family allowing their representatives to survive and flourish in Europe during the Miocene environmental transition. However, after having reached giant sizes and masses by the end of the Miocene, the extreme opening of the landscapes and the development of seasonal forests with deciduous leaves limiting the food supply (**Suc et al., 1999**; **Kovar-Eder, 2003**; **Jiménez-Moreno et al., 2010**) could have initiated the disappearance of the family.

## Acknowledgements

The authors express their gratitude to Loïc Costeur for giving them access to the collection of deinotheres hosted at the Natural History Museum of Basel and to Renaud Roch who produced copies of some of the specimens. We are grateful to Ursula Göhlich for providing the raw data of her recent publication. We also thank Davit Vasilyan for his help finding studies published in Russian. Finally we are also grateful to the two reviewers, Martin Pickford and an anonymous one, whose comments helped us to improve this manuscript.

## Author contributions

FG and DB conceived the study. FG performed the initial description, prepared the graphic figures, performed the first analysis and interpretations of the data and wrote an initial draft. OM took the pictures of the specimens, prepared illustration plates. DB and OM corrected the dataset, completed the interpretations and revised the manuscript.

## Data availability

Supplementary data are available on Zenodo (doi: 10.5281/zenodo.4468801).

## Supplementary information

- List of the European localities which yielded Deinotheriidae and the associated references. This dataset was used to construct **Figure 11** (doi: 10.5281/zenodo.4468801)

